# Pregnancy data enable identification of relevant biomarkers and a partial prognosis of autism at birth

**DOI:** 10.1101/2020.07.08.192989

**Authors:** Hugues Caly, Hamed Rabiei, Perrine Coste-Mazeau, Sebastien Hantz, Sophie Alain, Jean-Luc Eyraud, Thierry Chianea, Catherine Caly, David Makowski, Nouchine Hadjikhani, Eric Lemonnier, Yehezkel Ben-Ari

## Abstract

Attempts to extract early biomarkers and expedite detection of Autism Spectrum Disorder (ASD) have been centered on postnatal measures of babies at familial risk. Here, we suggest that it might be possible to do these tasks already at birth relying on ultrasound and biological measurements routinely collected from pregnant mothers and fetuses during gestation and birth. We performed a gradient boosting decision tree classification analysis in parallel with statistical tests on a population of babies with typical development or later diagnosed with ASD. By focusing on minimization of the false positive rate, the cross-validated specificity of the classifier reached to 96% with a sensitivity of 41% and a positive predictive value of 77%. Extracted biomarkers included sex, maternal familial history of auto-immune diseases, maternal immunization to CMV, IgG CMV level, timing of fetal rotation on head, femoral length in the 3rd trimester, white cells in the 3rd trimester, fetal heart rate during labour, newborn feeding and newborn’s temperature difference between birth and one day after. Statistical models revealed that 38% of babies later diagnosed with ASD had significantly larger fetal cephalic perimeter than age matched neurotypical babies, suggesting an in-utero origin of the bigger brains of toddlers with ASD. Results pave the way to use pregnancy follow-up measurements to provide an early prognosis of ASD and implement pre-symptomatic behavioral interventions to attenuate efficiently ASD developmental sequels.

## Introduction

Autism Spectrum Disorder (ASD) is characterized by persistent communication and social interactions deficits, and restricted, repetitive behaviors (DSM 5 -APA 2013) [1–3]. Since the first studies in the 1960s, its prevalence has steadily increased from 0.041% to 1.68% (CDC 2018) [4]. This increase is due to modifications of diagnostic criteria, wider access to diagnosis, and a genuine increase due to as yet undetermined factors, most likely a combination of genetic and environmental components [5,6]. In spite of the incidence of autism, there is yet no FDA or EMA approved drug agent to treat its core symptoms.

Clinical and histological observations are compatible with the notion that ASD is generated in the womb. Thus, increased ASD incidence has been related to maternal viral or microbial infection, with activation of the immune system [7–9], maternal influenza [9,10], drugs taken during pregnancy notably of sodium valproate [11], or exposure to environmental hazards [12,13]. Post-mortem analysis of brains from children with ASD reveals an abnormal excess of neurons in the prefrontal cortex indicative of an in-utero origin [14]. Brain overgrowth and megalencephalic brains have been reported in a subpopulation of children and adolescents with ASD [15,16] but whether this is initiated already in utero is controversial [17–21]. ASD is linked with many perinatal factors including emergency C-Section delivery, obstetric complications and preterm delivery reflecting the continuity between an in-utero insult parturition and birth [22].

Experimental data also suggest an in-utero pathogenesis of ASD [23–28]. Thus, maternal immune activation or valproate administration are associated with ASD [24,29] and early post-natal alterations are observed in genetic forms of autism [30]. In addition, brain overgrowth during parturition and birth has been observed in ASD animal models [29]. Therefore, determining the alterations occurring in-utero are instrumental in order to understand the pathogenesis of ASD.

Here, we reasoned that if ASD brain changes are already present during pregnancy, it might be possible to extract biomarkers from biological and imaging features that are routinely collected from the first pregnancy trimester to 1 day after birth, and give a prognosis of ASD shortly after birth. To this goal, we analyzed retrospectively those features in babies who went on being diagnosed 4-5 years later with ASD, and in a matched population of Neurotypical (NT) babies. Due to large number of features and complex multivariate and poorly understood links between them, we used several statistical tools to reveal patterns that distinguish NT babies from those with ASD.

A supervised machine learning (ML) algorithm was trained to classify babies in two groups, ASD and NT. A cross-validation (CV) technique was used to ensure the generalizability of the classifier’s results on an unseen future independent cohort. Features with highest impact on the classifier’s decisions were extracted and analyzed more precisely. In parallel, significant changes in distribution of all collected features between NT and ASD babies were identified through conventional statistical hypothesis tests. Finally, longitudinal developmental trajectories of fetuses were analyzed by statistical models to investigate the possibility that megalencephalic ASD brains in children and adolescents are generated in utero.

The use of follow-up features routinely collected in maternities without expensive additional tests will facilitate early behavioral treatments known to be more efficient when initiated before the end of the developmental plasticity period [31,32].

## Results

A classification algorithm with Shapley additive explanations (SHAP) framework in parallel to statistical hypothesis tests and statistical models were used to extract ASD biomarkers (see material and methods).

### Prognosis of ASD relying on the gradient boosting decision tree classifier

To classify babies in ASD and NT groups, a gradient boosting decision tree classifier was trained on collected features by adopting a strategy to minimize false ASD detections in the first place while keeping the true ASD detection rate as high as possible. The performance of the classifier with two feature selection strategies (FSS) was evaluated by the estimation of classification scores through averaging on 100 rounds of train-test coming from 10 times repeated 10-fold CV process.

In the semi-automatic FSS where a preselection of features based on domain knowledge was followed by Lasso regularization, the true negative rate (TNR aka specificity) i.e. the proportion of NT children correctly classified as NT, was of 0.96 (95% CI = [0.95, 0.97]), thus only 4% of NT children were wrongly classified as ASD (**Table 1**). The true positive rate (TPR aka sensitivity), i.e. the proportion of children with ASD correctly classified in the ASD group, was of 0.41 (95% CI = [0.37, 0.45]). However, the positive predictive value (PPV aka precision) was as high as 0.77 (95%CI = [0.72, 0.81]), implying that 77% of babies classified as ASD were indeed diagnosed later as children with ASD. Therefore, NT children were almost completely correctly identified at birth and a prognostic of ASD could be made in a subgroup of children with a high precision.

**Table 1.**
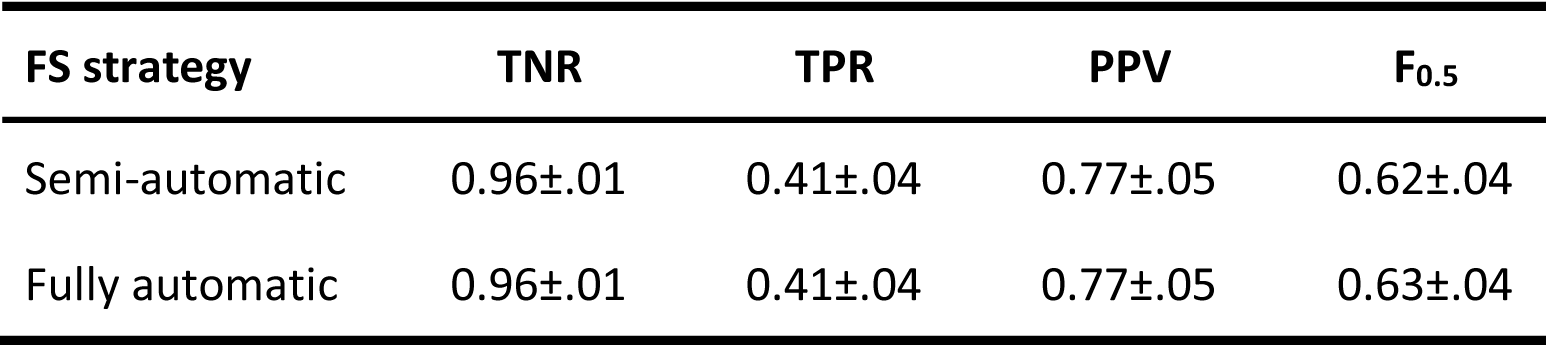
Evaluation of the classifier performance. Estimated classification scores with 95% confidence intervals computed through averaging on a cross-validation process based on two feature selection (FS) strategies. TNR true negative rate, TPR true positive rate, PPV positive predictive value.

If we let the classifier to select features automatically without any medical presumptions, the classifier achieves the same performance as in the semi-automatic FSS (**Table 1**). It shows the efficiency of the classifier to cope with a large feature space at least as good as when the feature space was pruned by medical presumptions.

### Extraction of important features

To extract features that play an important role in the classification process, we considered two approaches.

### Feature frequency in CV folds

In the CV process, the classifier is trained from scratch in each fold and selects features that distinguish better NT babies from ASD ones existing in the training set. Features that have been selected by the classifier in at least half of the 100 CV folds are given in **Fig. 1**. Fetal rotation on head, Femoral length percentile in the 3^rd^ trimester (T3), Cephalic perimeter percentile in the 2^nd^ trimester (T2), Breast feeding, Sex, Ratio of cephalic perimeter to femoral length in T3 and Fetal heart rate during labour (FIGO classification) have been selected in both FSSs.

**Figure 1.**
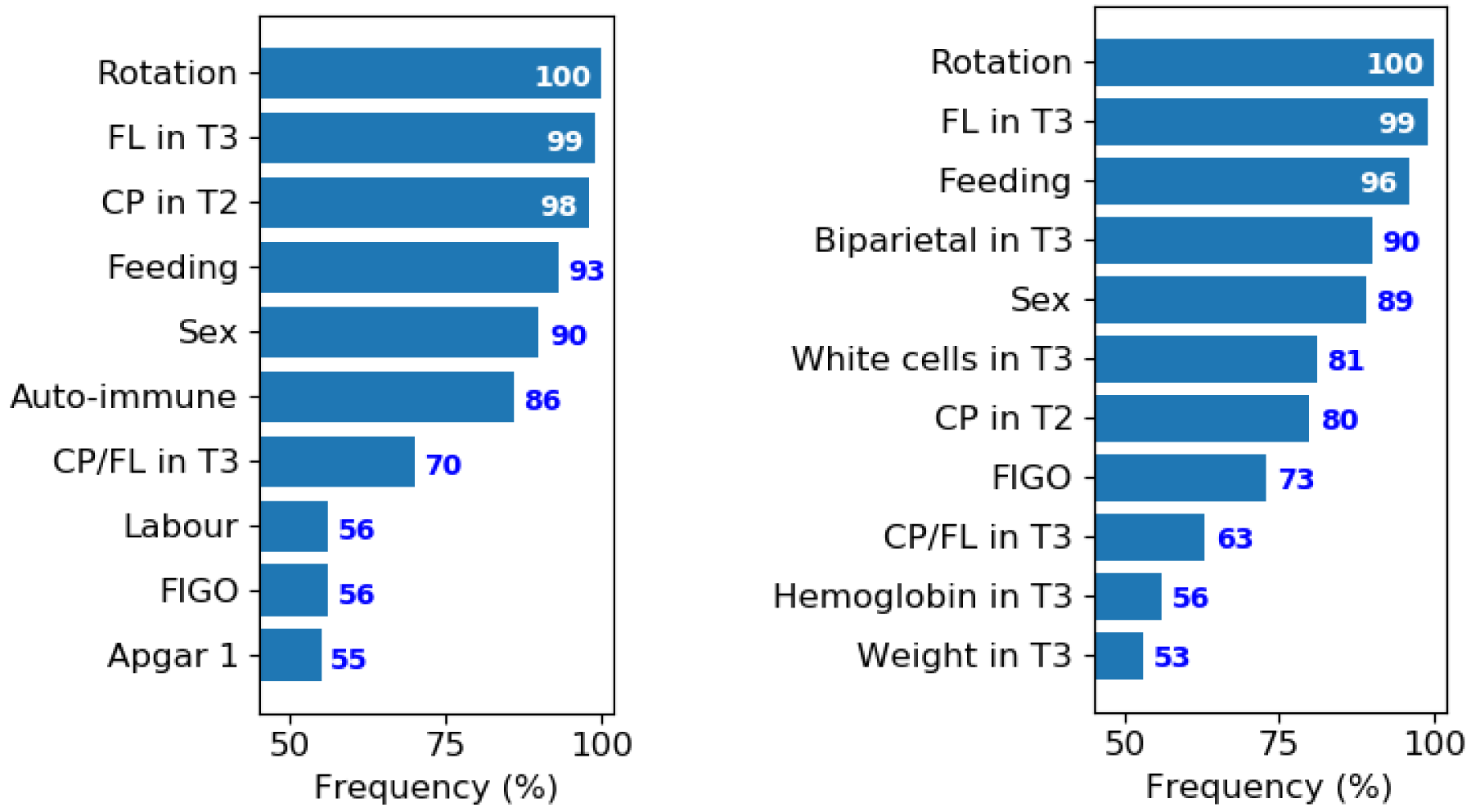
Extraction of biomarkers through feature frequency in the cross-validation process. Features that have been selected by the semi-automatic (left plot) and fully automatic (right plot) feature selection strategies in more than 50 out of 100 folds of the CV process. Features with higher frequencies are more important to the classifier which means they separate better NT babies from ASD ones. T2, second trimester. T3, third trimester. Rotation, Fetal rotation on head (days). FL, Femoral length percentile. CP, cephalic perimeter percentile. Feeding, breast feeding. Auto-immune, familial maternal history of auto-immune diseases. CP/FL, ratio of cephalic perimeter to femoral length. Labour, duration of the first part of the labour. FIGO, fetal heart rate during labour (FIGO classification). Apgar 1, Apgar score in 1 minute. Biparietal, biparietal diameter percentile. Weight, fetal weight percentile estimation.

In the semi-automatic FSS, Familial maternal history of auto-immune diseases, Duration of the first part of the labour and Apgar score in 1 minute also appeared frequently. On the other hand, Biparietal diameter in T3, White cells in T3, Hemoglobin in T3 and Fetal weight estimation in T3 were also selected as important features by the automatic FSS while they were considered as medically irrelevant. In other words, the classifier with automatic FSS could detect patterns in some features that are not normally considered relevant to ASD.

Some features were selected with less frequency and many features have never been selected since they were considered as irrelevant by the classifier. A complete list of features that have been selected by the classifier at least once in the CV process is given (**Supplementary Table S1)**.

### Feature impacts

Our second approach to extract important features relies on SHAP framework. It computes the impact of each feature on the classifier’s output and also determines a range of values of the feature that increase the probability of babies to be classified as NT or ASD. SHAP values of 5 features with the highest relative impact are shown in **Fig. 2** for all babies and for both feature selection strategies. In each feature line, a point colored by the corresponding feature value represents one baby and the color map indicates how each feature’s impact varies according to its values. Feature values situated in the positive or negative SHAP side (orange or green regions) leads to ASD or NT classification, respectively. The relative impact of these features together with their range of values that increase the probability of ASD classification are given in **Table 2**.

**Table 2.** Features with the highest impact on classifier based on SHAP analysis. For each feature, the relative impact and the alarming range i.e. the range of values that push the classifier to ASD decision are presented. T3, third trimester. Rotation, Fetal rotation on head. FL, Femoral length percentile. Feeding, Breast feeding. Auto-immune, Familial maternal history of auto-immune diseases.

| A. Semi-automatic feature selection strategy |  |  |
| --- | --- | --- |
| Feature | Relative impact | Alarming range |
| Rotation | 52% | Earlier than 148 days |
| FL in T3 | 19% | Higher than 72% |
| Feeding | 9% | Mixed of Maternal and artificial |
| Sex | 7% | Male |
| Auto-immune | 4% | Yes |
| B. Fully automatic feature selection strategy |  |  |
| Feature | Relative impact | Alarming range |
| Rotation | 44% | Earlier than 148 days |
| White cells in T3 | 16% | Less than 9100 |
| FL in T3 | 13% | Higher than 72% |
| Sex | 9% | Male |
| Feeding | 5% | Mixed of Maternal and artificial |

**Figure 2.**
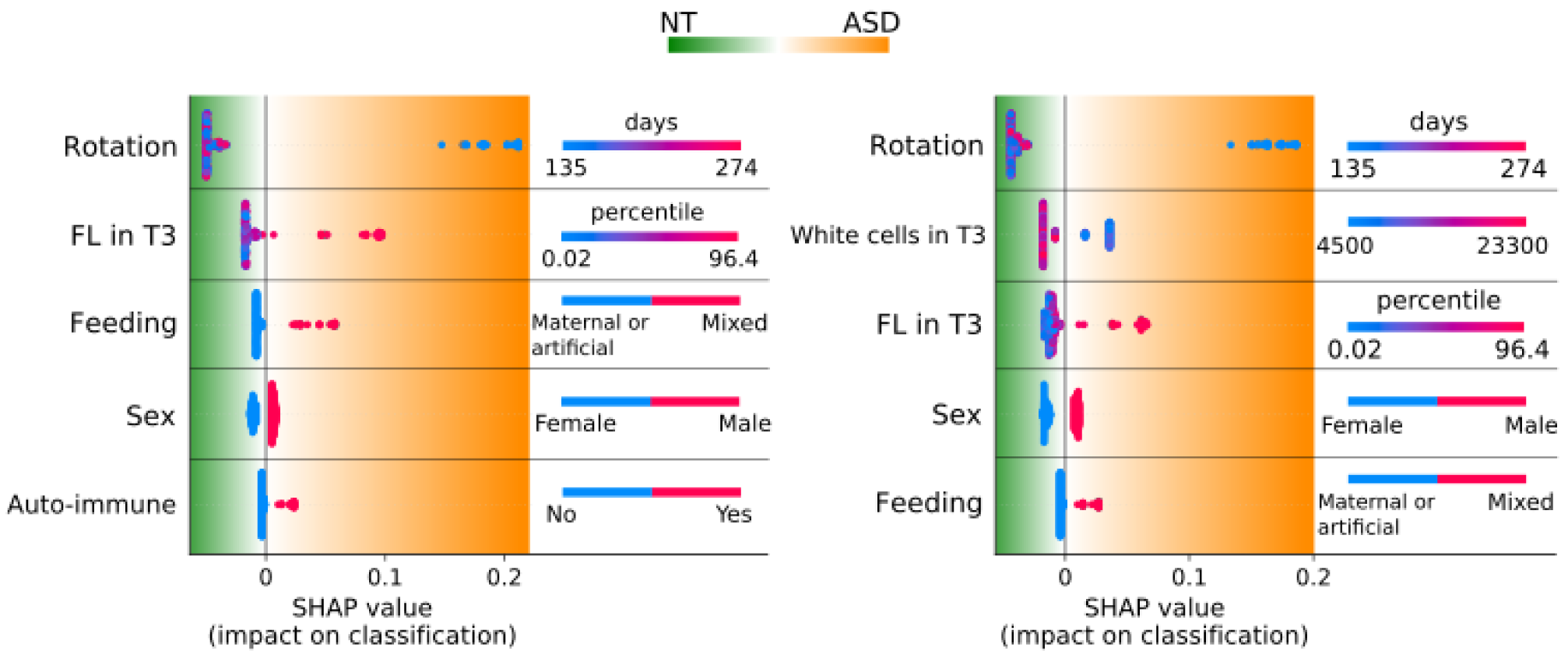
Features with the highest impact on the classifier based on SHAP analysis. Left: classification with semi-automatic FSS. Right: classification with fully automatic FSS. The impact (SHAP values) of features on the classification as NT or ASD is shown. Features are ordered from top to down according to their decreasing impact. Each point at each feature line represents a baby colored by the corresponding feature value. In each plot, feature values that lead the classifier to an NT or ASD prognosis are on the green and orange regions, respectively. T3, third trimester. Rotation, Fetal rotation on head. Feeding, Breast feeding. FL, Femoral length percentile. Auto-immune, Familial maternal history of auto-immune diseases.

Thus, in both feature selection strategies, fetuses who rotated on head before 148 days of gestational age were more likely to be classified in the ASD group. In fact, statistical analysis shows 35.09% of ASD babies rotated earlier than 148 days, which is significantly different from 3.72% of NT ones who rotated in that period (*χ* ^2^(1, N=245) = 40.74, p<0.001). High values of femoral length percentile in T3 (higher than 72%) led to an ASD prognostic by the classifier. The proportion of ASD babies with femoral length percentile larger than 72% in T3 is 24.56% whereas 7.94% of NT babies have a large femoral length in this range (*χ* ^2^(1, N=246) = 10.10, p=0.001).

Feeding babies with a mixture of maternal and artificial milks led to ASD classification. 17.86% of ASD babies were fed in a mixed way while the proportion is 5.91% in NT group (*χ* ^2^(1, N=242) = 6.31, p=0.01). Boys were more likely to be classified as ASD than girls. In ASD group, 80.95% of babies are male whereas the proportion of males in NT group is almost balanced with 48.68% (*χ* ^2^(1, N=252) = 18.76, p<0.001).

Familial maternal history of auto-immune diseases was considered as an important feature by the semi-automatic FSS. The proportion of ASD babies with familial maternal history of auto-immune diseases is about 19.05% whereas the proportion of NT babies with this feature is 6.35% (*χ* ^2^(1, N=252) = 4.73, p=0.006). For the fully FSS, white cells less than 9100 in T3 led the classifier to ASD decision. Among ASD babies, 47.54% of them have white cells less than 9100 whereas the proportion of NT babies is about 27.65% (*χ* ^2^(1, N=231) = 7.17, p=0.007). A complete list of features with nonzero relative impact is given in **Supplementary Table S1**.

In summary, classification process helped to extract specific prognostic biomarkers among a lot of recorded features, and SHAP analysis revealed patterns in those biomarkers that are significantly different in NT and ASD groups.

### Statistical difference in feature distributions between ASD and NT

Independent from the classification process, we ran statistical hypothesis tests on all recorded features. Results for features with significantly different distributions in NT and ASD groups are given in **Table 3 and Fig. 3** (see also **Supplementary Table S2** for all other features). In case of categorical features, number (n) and frequency (%) of babies in each group and the results of Chi-square test (*χ* ^2^) are given. For the numerical feature, number of samples, median, mean, standard error of mean (SEM) and 95% confidence interval of mean of feature values in each group together with results of Mann-Whitney U test (MWU) are presented.

**Table 3.** Results of statistical tests for features which are significantly different between NT and ASD groups. n, number of samples. SEM, standard error of mean. CI, confidence interval. *χ* ^2^, Chi square test. MWU, statistics of Mann-Whitney U test.

| Feature |  | Statistics |  |  |
| --- | --- | --- | --- | --- |
|  |  | NT group | ASD group | Test results |
| Sex | Male % (n) | 48.68 (92) | 80.95 (51) | $\chi^2(1, 252) = 18.76$<br>p<0.001 |
|  | Female % (n) | 51.32 (97) | 19.05 (12) |  |
| Absolute child's temperature difference day 1 – birth | > 1°C % (n) | 14.00 (21) | 41.67 (20) | $\chi^2(1, 198) = 15.31$<br>p<0.001 |
|  | < 1°C % (n) | 86.00 (129) | 58.33 (28) |  |
| CMV | Immunized % (n) | 36.62 (26) | 76.92 (20) | $\chi^2(1, 97) = 10.83$<br>p<0.001 |
|  | Negative % (n) | 63.38 (45) | 23.08 (6) |  |
| Fetal heart rate during labour (FIGO classification) | Pathological % (n) | 10.20 (15) | 28.89 (13) | $\chi^2(2, 192) = 13.82$<br>p = 0.001 |
|  | Suspect % (n) | 28.57 (42) | 8.89 (4) |  |
|  | Normal % (n) | 61.22 (90) | 62.22 (28) |  |
| IgG (IU) | n | 64 | 22 | MWU = 382.5<br>p<0.001 |
|  | Median | 0 | 9950 |  |
|  | Mean | 4675 | 10913.64 |  |
|  | SEM | 963.19 | 2034 |  |
|  | 95% CI | [2750.23, 6599.77] | [6683.70, 15143.57] |  |

**Figure 3.**
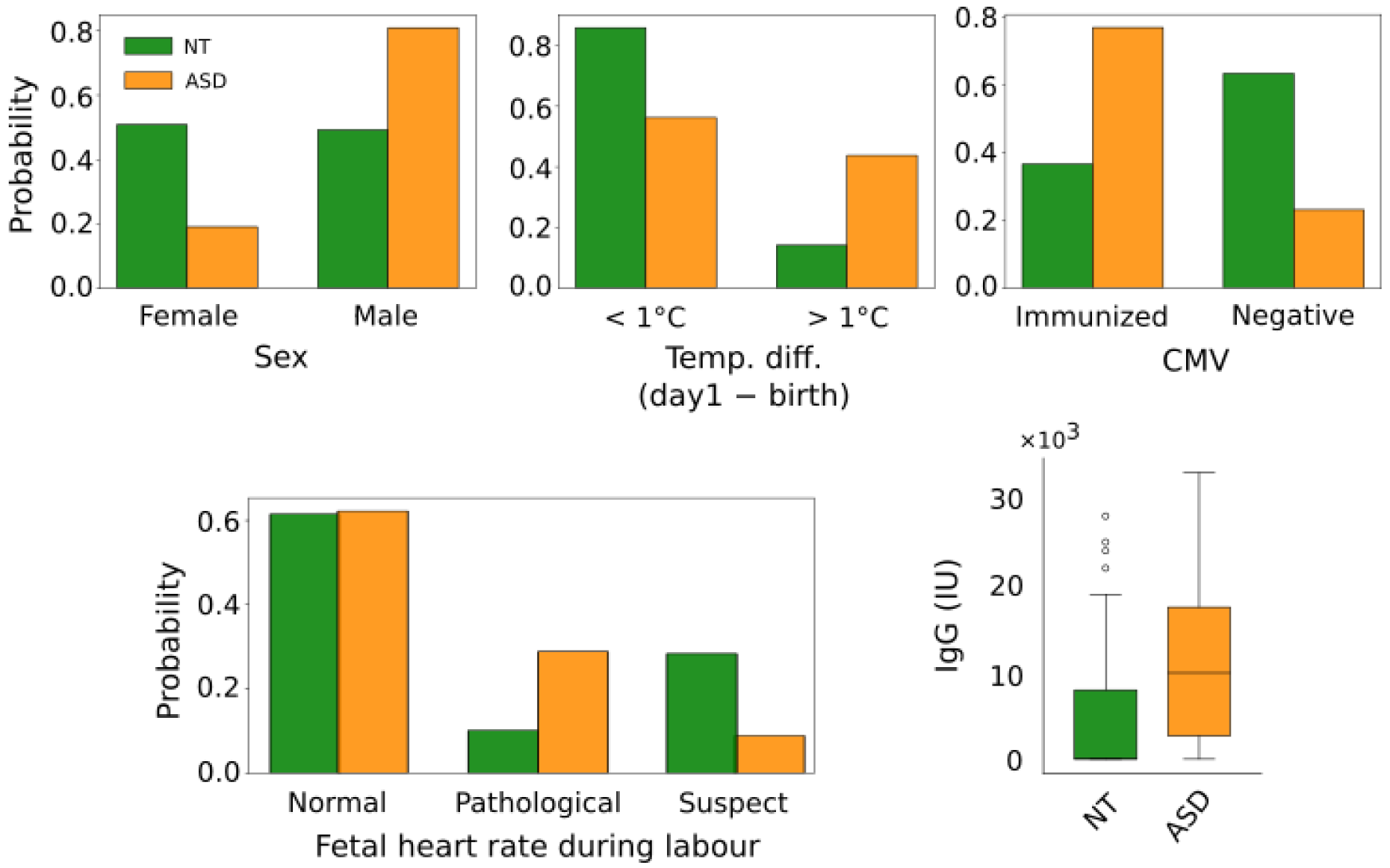
Normalized distribution of statistically significant features. Distributions of Sex, Absolute child’s temperature difference between birth and 1 day later, CMV immunoreactivity, Fetal heart rate during labour (FIGO classification) and IgG levels are shown in NT (green) and ASD (orange) groups. In the boxplot center line is median, box limits are upper and lower quartiles and whiskers are 1.5x interquartile range. See **Table 3** for quantitative comparisons.

Among NT children, 48.68% are male versus 80.95% of ASD (*χ* ^2^(1, 252) = 18.76, p<0.001). 14% and 41.67% of NT and ASD newborns respectively had a temperature difference of more than 1°C (in either direction) between birth and day 1 (*χ* ^2^(1, 198) = 15.31, p<0.001) (see also **Supplementary Fig. S1**). With cytomegalovirus serology (CMV), 36.62% and 76.92% of NT and ASD mothers were immunized respectively (*χ* ^2^(1, 97) = 10.83, p<0.001). Blood samples used for Guthrie’s test confirmed the lack of congenital hypothyroidy, mucovisidosis, drepanocytosis, phenylketonuria and congenital adrenal gland hyperplasia. They also revealed no CMV mRNA indicating that, with the limits of this test, the impact is not due to neonatal viral infection but to maternal immunization. The median of IgG CMV is 0 IU in NT children versus 9950 IU in the ASD group (MWU = 382.5, p<0.001). The strong difference of the median (0) and mean (4675.00) in the NT group reflects a right skewness of the IgG curve with at least 50% of NT children having 0 IgG CMV levels (**Fig. 3**). With the FIGO classification of fetal heart rate during labour, 61.22% and 62.22% of NT and ASD children respectively have a normal heart rate whereas 10.20% and 28.89% of NT and ASD children respectively have a pathological heart rate (*χ* ^2^(2, 192) = 13.82, p = 0.001).

### Cephalic perimeter growth differs in NT and ASD

Regression analysis shows that during the 2^nd^ trimester, the cephalic perimeter (CP) growth rate is significantly different between NT and ASD (ANCOVA, p=0.046) with slopes of regression line equals to 1.35 (p<0.001, R^2^=0.52, ρ=0.72 Pearson’s correlation coefficient) for NT and 1.73 (p<0.001, R^2^=0.61, ρ=0.78) for ASD (**Fig. 4a**). However, the mean percentile values of CP are not different in this age (t-test, p=0.2) (box plot of **Fig. 4a**). During the 3^rd^ trimester, the increasing growth slopes are similar for NT (1.11, p<0.001, R^2^=0.23, ρ=0.48) and for ASD (1.01, p<0.001, R^2^=0.20, ρ=0.45) (ANCOVA, p=0.72), although there is an increasing trend in the CP percentile (MWU, p=0.83). In contrast, before birth, ASD’s CP percentiles are significantly higher than NT (MWU, p=0.02), but the growth slopes are similar as 0.65 (p<0.001, R^2^=0.46, ρ=0.68) for NT and 0.63 (p=0.01, R^2^=0.20, ρ=0.45) for ASD (ANCOVA, p=0.93; see also **Supplementary Fig. S2**). The quadratic mixed effect model shows a similar (p=0.30) slowdown of CP increase in the NT and ASD groups along the gestation (**Fig. 4b**). The coefficient of quadratic term is -0.006 for NT (95% CI: [-0.006, -0.005]; p<0.001) and -0.005 for ASD (95% CI: [-0.006, -0.004]; p<0.001). The larger CI of the ASD group suggests a larger heterogeneity in ASD than in NT.

**Figure 4:**
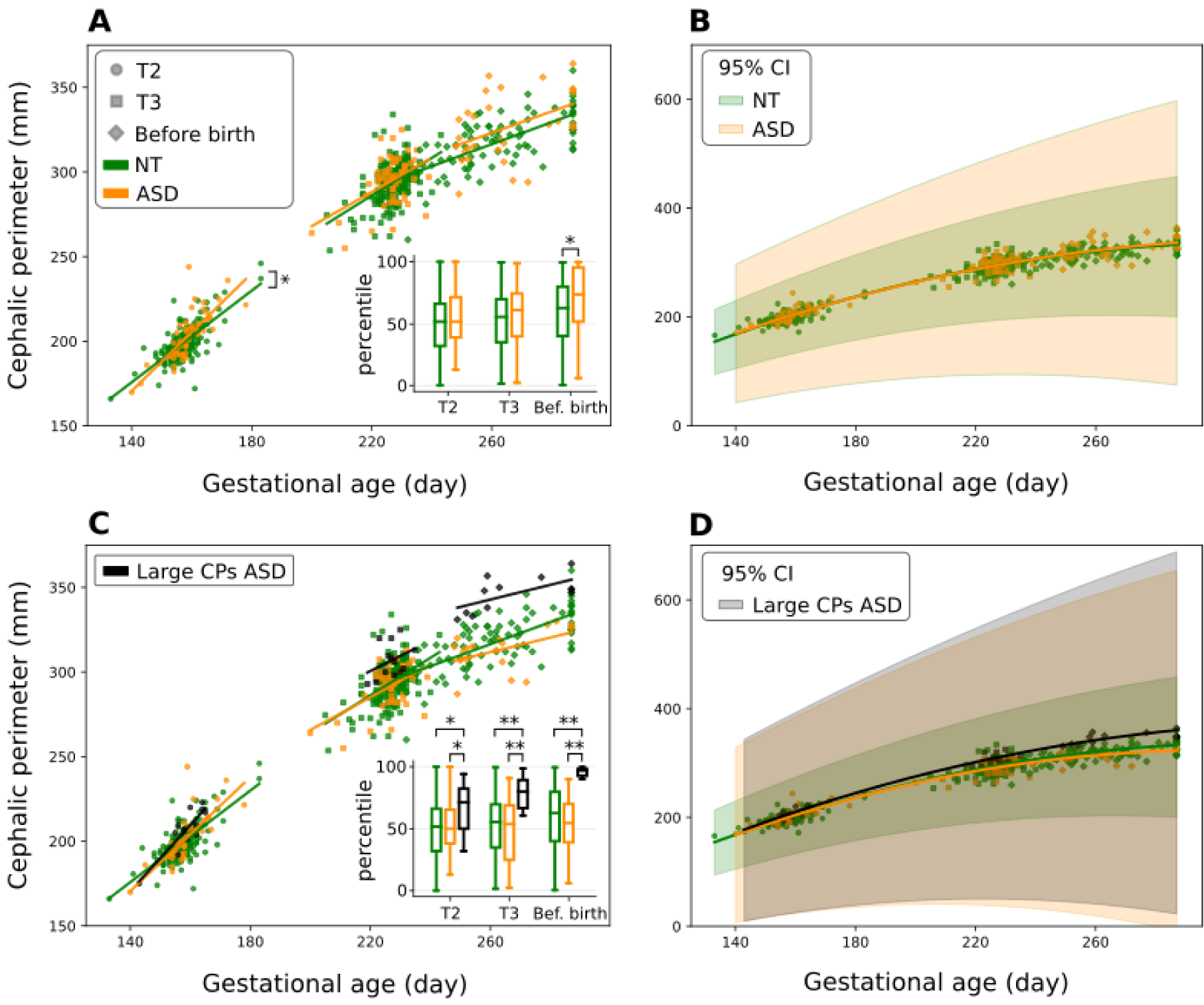
Cephalic perimeter (CP) growth is slowed down during development in NT and ASD, but shortly before birth ASD CP is bigger than NT CP. The CP in NT (green) and ASD (orange) groups is depicted versus the gestational age in T2 (circle), T3 (square) and shortly before birth (diamond) periods. **A)** Linear regression analysis shows the growth of CP in T2 is significantly higher in ASD than NT (p<0.05). Also, ASD CPs are bigger than NT shortly before birth (p<0.05) as shown in the boxplot (center line, median; box limits, upper and lower quartiles; whiskers, 1.5x interquartile range). **B)** The quadratic mixed effect model shows similar progressive slowdown of CP of NT and ASD children towards birth. 95% confidence intervals are shown for each curve. **C)** A subpopulation of ASD group with large CPs before birth was separated from the rest of the ASD group (black points). Linear regression analysis shows similar CP growth rate in the 3 groups. However, ASD embryos with large CPs have significantly larger CPs than NTs in the 2^nd^ and 3^rd^ trimester and shortly before birth (p<0.05, p<0.001 and p< 0.001, respectively). **D)** The quadratic mixed effect model shows similar progressive slowdown of CP of NT, ASD and “Large CPs ASD” children towards birth. * p<0.05, ** p<0.001

Exploration of CP percentile distributions revealed that 38% of ASD babies have CP percentile higher than 90% before birth (box plot of **Fig. 4a** and **Supplementary Fig. S2**). We separated those ASD babies into a new group called “Large CPs ASD” (**Fig. 4c**) to investigate whether they have already had larger CPs in T2 and T3. We found that during the 2^nd^ trimester, there is a significant difference in CP percentile between groups (ANOVA, p=0.02) with children in “Large CPs ASD” being bigger than both NT (Tukey, p=0.01) and the remaining ASD subpopulation (Tukey, p=0.04) (boxplot of **Fig. 4c**). The significant difference is also observed during the 3^rd^ trimester and before birth (Kruskal-Wallis, p<0.001) where “Large CPs ASD” have larger CPs than both NT and remaining ASD subpopulation (Dunn, p<0.001 for both groups).

Linear regression analysis with the ANCOVA test shows no difference in the CP growth rate between the 3 groups in either period (**Fig. 4c**). The decline in head growth rate during gestation was confirmed in “Large CPs ASD” like for the other groups with the quadratic mixed effect model (**Fig. 4d**) with a coefficient of quadratic terms of -0.006 in ASD (95% CI: [-0.008, -0.005] ; p<0.001) and -0.005 in “Large CPs ASD” (95% CI: [-0.006, -0.004]; p<0.001).

Therefore, the children of the ASD group with large CPs before birth have already had large CPs in the 2^nd^ and 3^rd^ trimesters. Moreover, a slowdown of CP growth similar to that of NT and other ASD children was observed in the ASD group with large CPs. Results suggest a bigger heterogeneity of CP growth in ASD that however preserves the progressive CP attenuated growth in preparation for birth as in naïve brains.

## Discussion

The difficulty of developing an early diagnostic of ASD stems from its prenatal and early postnatal generation and the heterogeneity of its symptoms with an impairment of motor skills, visual perception, social interactions or attention to faces, that all take time until they become noticeable. The inaugurating insult leading to ASD alters cell proliferation, migration, synapse formation, pruning and formation of functional cell assemblies. This cascade of impairments leads to the classical social deficits and repetitive behaviors that typically emerge around 24 months of age, and to heightened sensitivity to stimuli from many modalities. As early behavioral treatment ameliorates ASD deficits and attenuates long-term outcomes [31,32], early detection in toddlers is essential before clinical signs are conspicuous.

Several attempts have been made to detect ASD early relying on neuroimaging techniques, EEG measures or genetic variants. In these studies, the prediction is centered primarily on siblings of children diagnosed with ASD, that is, high-risk populations. They are therefore hampered by this factor, as the ratio of high-risk to low-risk is not representative of the general population.

Neuroimaging in babies at high familial risk of autism have revealed increased brain volume that appears before ASD diagnosis [33–35]. The authors obtained a high sensitivity and accuracy of ASD prediction, but the restriction to high-risk sibling hampers and limits the generalizability of the conclusion to first born without siblings with ASD [36]. Similarly, EEG power spectrum analysis of at-risk siblings from three months onwards [37] distinguishes ASD from NT children with an accuracy (true negative and positive outcome) of 91.67%. The positive predictive value is, however, about 63.93% of those diagnosed as at risk during the first year go on to develop ASD later. Interestingly, the frontal EEG at age 3-12 months most accurately discriminated the ASD group, pointing to early perinatal processes vs. later ones, and the presence of early subclinical changes that can be detected by early frontal EEG power. The lower gamma power observed is suggestive of an imbalance between excitation and inhibition. The widely used genocentric approach has not allowed establishing an early prognostic of ASD to large populations due to several limitations. Hundreds of genetic mutations and variants have been identified often with poor penetration that produces incremental risks when cumulated [38–43]. In addition, *de novo* variations play an important role [44], complicating the prediction. Moreover, non-genetic factors play an important role in the pathogenesis including environmental factors during maternity (e.g. pollution, pesticides, prenatal vitamins – for review [13]) are instrumental in ASD pathogenesis as they augment the incidence of ASD. However, they cannot provide an early prediction of ASD.

Our goal here was to determine whether it is possible to extract prognostic biomarkers associated with ASD from imaging and biological features that are routinely collected during pregnancy and birth, and give a prognosis of ASD shortly after birth. We reasoned that this would on one hand provide compelling evidence that ASD is born in the womb, and on the other hand offer a wide range of novel possibilities by using data normally available in maternity wards. In this aim, ML algorithms and conventional statistical hypothesis tests were employed to analyze data collected from a representative population of ASD with a global incidence (1.21%) similar to that reported in Europe and other countries. ML is useful in this context, as it enables to identify features that are poorly or not statistically significant, but converge to impact ASD identification. Moreover, ML approaches have shown recently their power in disease prognosis with applications in e.g. hepatitis prediction [45], classification of diabetic patients [46,47] and lung cancer screening [48]. They have also recently enabled to give brain specific interactions probability of each gene with all the genes of the network and their probability association with ASD [49] but without differentiating NT and ASD babies.

Results suggest that a combination of collected features intuitively linked to ASD and others not associated with ASD impact the classification and prognosis. Many of these have, at this stage, no straightforward mechanistic links with ASD, except quite indirect speculative connections. The femoral length percentile differences might be related to the finger and toe ratios altered in ASD because of hormonal influences [50,51]. Gestational hypoxia [52] like pathological heart rate during labor and birth has been associated with neurological sequels [53]. 95% of embryos have their head down at birth [54,55], but here we show that the shift occurs earlier in ASD possibly suggesting an earlier preparation for birth. There are less than 1°C changes in body temperature in the majority of NT children between birth and 1 day later, but bigger differences (warmer or cooler) in ASD. This suggests a difficulty in controlling body temperature that might be related to inflammatory signals [56]. Several features associated with inflammatory signals are also significantly different in ASD and NT, including maternal immunization to CMV, the average of IgG CMV units, and familial maternal history of auto-immune diseases [9]. Other impacting parameters such as low values of White cells in T3 and mixed maternal and artificial milk do not have documented links with ASD.

The developmental curve of Cephalic Perimeter (CP) in utero suggests that brain growth is impacted at a very early stage (also see [17]). Brain growth of NT and ASD is slowed down from the 2nd trimester to birth but with important differences between them. Although the mean CP values are not different between NT and ASD, there is a significant acceleration of growth in the latter versus the former in the 2^nd^ trimester suggesting a long-lasting impact of the pathogenic event such that ASD group has significantly larger CP before birth. We also identified a subpopulation of “Large CPs ASD” with significantly larger CPs than age matched NTs during the 2^nd^ and 3^rd^ trimesters and before birth. Interestingly, the CP of a subpopulation (15%) of children and adolescents with ASD has been reported to be bigger than NT with “megalencephalic” features [15,16]. Therefore, brain growth process is impacted already from the 2^nd^ trimester with a CP that continues growing during the few days that precede birth. Future studies will have to determine if the brain continuous to grow *during* parturition as observed in animal models (see below).

Experimental observations are in accord with this. Hippocampal and neocortical volumes are increased in an animal model of ASD and hippocampal neuron size grows during parturition and birth [25,29]. Neurons with immature features are present in the adolescent and adult human amygdala [57]. In patients with ASD, the process that governs postnatal cellular maturation, like the trajectory of neuronal development, is altered in the human amygdala with a persistence of neurons endowed with immature features [58,59]. This “immaturity” of impacted neurons stands at the core of the “neuroarchaeology” concept positing that the inaugurating insult in utero deviates developmental sequences leading to misplaced or misconnected neurons that present signs of immaturity and generate patterns that disturb behaviorally relevant oscillations [60]. Neurons with immature properties have been observed in many pathological conditions in experimental models of ASD. High (Cl-)i and GABA excitatory actions are observed in immature neurons in MIA [25], maternal Valproate and Fragile X [30,61], and Rett syndrome [62] suggesting common global reaction to the pathogenic insult. Restoration of GABAergic inhibition also attenuates in humans and rodents the severity of ASD paving the way to novel therapeutic strategies based on selective actions on immature neurons [63,64].

There are many limitations to the present study. The small sample size and the small number of girls limit the generalizability of the results. On the other hand, according to results of statistical tests, some features such as child’s temperature, CMV and IgG are distinguishing but were not included in the classification process due to their missing values. Moreover, we deliberately preferred to minimize false positives, which restricted ASD detection rate and feature extraction. Therefore, these results are not meant to provide an early diagnosis of ASD, but a possible prognostic tool and a proof of concept. Future studies might help ameliorating these aspects by considering a larger population in order to cope with the heterogeneity of ASD features.

To conclude, our results suggest that it might be possible to establish a prognosis at birth of a subpopulation of babies who will develop ASD. The trained algorithm will require larger replications before being considered as a clinical tool for predicting ASD in large populations, as false predictions might adversely affect individuals. Yet, results in keeping with large evidence suggests that ASD is generated by a pathogenic sequence of events in-utero, that impacts essential developmental processes from cell proliferation and migration to neuronal growth, synapse formation, and network construction [41,60]. The time and structural basis of the inaugurating insult most likely underlies the heterogeneity of ASD [41,60]. If confirmed, the identification at birth of babies at risk of ASD relying on data that are routinely available in maternity wards, without additional techniques, will facilitate the use of behavioral preventing therapeutic strategies before the end of the developmental plasticity critical period [31,32]. Our approach might also be completed by frontal EEG or genetic data in order to improve the accuracy and sensitivity of the prediction.

## Methods

### Data and experiments

In 2012-2013, 5356 children were born in the maternity Hospital of the University of Limoges in France. Two to 5 years later, 65 of these children (1.21%) were diagnosed with ASD (DSM-5 criteria American Psychiatry Association 2013) and confirmed by ADI-R and Autism Diagnostic Observation Schedule (ADOS G). Two children were excluded (Trisomy 21 and extreme prematurity, birth at 30 weeks). ASD incidence rate in our population (1.21%) is close to the reported rate in the literature (CDC 2018), which justifies our sampling approach.

The files of the 63 children (12 girls and 51 boys) were matched with 189 Neurotypical (NT) children based on mother’s age, parity and term of childbirth. In following, for simplification, we shall refer to babies diagnosed years later as NT or ASD as NT or ASD babies respectively. For each mother and baby, 116 features were recorded during pregnancy until 1 day after birth. The feature space consists of 77 numerical features (e.g. mother’s BMI, ultrasound measurements), 38 categorical (e.g. gender, familial medical history, auditory tests), and 1 ordinal (placenta Grannum classification in the 3^rd^ trimester), which are commonly recorded in French maternity hospitals. **Supplementary Table S3** provides the entire list of features used in this study.

The goal of this study is to find patterns in recorded features that distinguish ASD babies from NT ones, and our approach to this goal is 2-folded. First, a supervised classification algorithm is trained on data and features with high impact on the classifier are extracted with two different methods. In the second approach, appropriate statistical hypothesis tests are performed to find features that have significantly different distributions in NT and ASD groups. Moreover, developmental trajectories of cephalic perimeter were studied by statistical models.

### Data preprocessing

The values of each 116 recorded feature in the dataset were explored and cleaned carefully to reduce the noise in computations. Features with missing value rate higher than 10% were removed from classification process to reduce the imputation bias in results. Features that were included in the classification process are given in **Supplementary Table S4**. Consequently, the classification dataset consists of 67 features for which 2.58% of values are missing in total. The one-hot encoding technique [65] was applied to binarize categorical features. To avoid co-linearity, one category of each feature was dropped.

### Feature selection for classification

A common issue in technology-based biological classification studies is the low ratio of sample size to number of collected features [66] which increases the classification error and the risk of data overfitting [67–69]. To treat this issue, a good practice is reducing the dimension of feature space by finding and dropping irrelevant and redundant features based on some criteria or domain knowledge.

In this study, features were selected by two strategies: fully automatic strategy by the Lasso regularization technique [70], and semi-automatic strategy which consists of a feature preselection based on medical knowledge followed by the Lasso technique. The goal was to compare classification performance with and without human intervention in feature selection and also, the similarity between selected features by those strategies.

The automatic strategy relies on a Lasso regularization technique that is applied directly in the training process of the classifier. It shrinks the impact of irrelevant features on classification and selects implicitly the most distinguishing ones. This technique is known to be very effective even in presence of many very irrelevant features [70] and may find some features that are not already known to be linked to ASD. The semi-automatic strategy investigated the effect of feature preselection by using domain knowledge before applying the Lasso technique. In this strategy, 19 features (out of 67) that might be linked to ASD were preselected and fed to training process (**Supplementary Table S4**). Among those features, the most informative ones were selected by the Lasso technique.

### Classification process

To classify children as NT or ASD, we used a model based on the gradient boosting decision tree algorithm [71]. This is a nonparametric supervised learning method which uses a tree-like model to infer a decision for each baby from feature values. Instead of using only one tree model, an ensemble of them is considered under the gradient boosting technique to fortify the ability of the classifier. Starting with a simple classification tree model, the model learns by adding more trees in an iterative manner to minimize a learning objective. It can detect complex underlying patterns of features to predict the binary target variable of belonging to the ASD group. This algorithm gives state-of-the-art results in a wide range of classification applications, especially in healthcare and diagnosis of diseases [45,46,72,73].

To implement the gradient boosting decision tree algorithm efficiently, we relied on the eXtreme Gradient Boosting (XGBoost) library [74]. Tuning its hyper-parameters to control the implementation of the algorithm enabled to resolve many classification problems (see https://github.com/dmlc/xgboost/blob/master/demo/README.md). Moreover, XGBoost has a built-in strategy to deal with missing values by finding the best imputation [74].

In this study, we used a nested cross-validation process to tune the number of decision trees and evaluate the classifier’s performance (see below). Moreover, we tuned carefully several hyperparameters to control the complexity of the model and avoid overfitting. Namely, the depth of each tree was set to a relatively low value as 5. The model weights were shrunk after each learning iteration by a factor of 0.01. Features were subsampled in each tree to make the model robust to potential noise in data. Rate of subsampling was inversely proportional to number of features and was selected as 0.7 and 0.5 in semi-automatic and fully automatic feature selection strategies (FSS) respectively. We used Lasso and Ridge regularization techniques to impose a penalty on the complexity of the classifier. Lasso regularization, as explained above, also helped to detect and remove less relevant features automatically which, in turn, avoided overfitting.

Imbalanced datasets are common in medical studies due to low prevalence of diseases. This causes a classifier to learn mostly patterns in the majority class, i.e. control samples. To cope with this issue, we imposed a higher weight on the misclassification error of ASD samples than that of NT ones. The classifier output for each baby is the probability of the baby to belong to the ASD group. We set the decision threshold to 0.5 to binarize the predicted probability.

By choosing higher weights on ASD misclassification error or lower values of the decision threshold than those considered here, more ASD children could be detected, but false positive rate would increase as well, which is in contrast to our ethical concerns and would increase the risk of overfitting. The list of XGBoost hyperparameters with corresponding values used in this study is given in **Table 4**.

**Table 4.** Hyperparameters of the XGBoost classifier. Feature subsampling rate is set to 0.7 and 0.5 for semi-automatic and fully automatic feature selection strategies respectively.

| Hyperparameter | Values | Hyperparameter | Values |
| --- | --- | --- | --- |
| Number of trees | 10, 20, 30 | Feature subsampling rate | 0.7 or 0.5 |
| Tree depth | 5 | ASD sample weight | 2 |
| Learning rate | 0.01 | Lasso & Ridge coefficients | 10 |

### Hyperparameter tuning and classifier evaluation

To tune hyperparameters of the XGBoost classifier and to evaluate its performance, we used a 10-times repeated nested 10-fold stratified cross-validation (CV) process. In each repetition of the CV process, the whole dataset was divided randomly in 10 partitions, 9 partitions for training the classifier and 1 held-out partition to test the trained classifier and to ensure that the algorithm can be generalized in future unseen samples. The train-test process of the classifier ran in 10 rounds. In each round, the hyperparameters of the classifier were tuned on train data through an internal 5-fold stratified cross-validation grid search on values given in the **Table 4**. Optimal hyperparameters were chosen to maximize *F*_*0*.*5*_ score of classification (see below for definition of *F*_*0*.*5*_ score). The model was trained using optimal hyperparameters on the train data and the trained model was used to predict the target variable of samples in the 1 held-out test partition. Beyond *F*_*0*.*5*_, local classification scores, including True Positive Rate (TPR aka sensitivity),

True Negative Rate (TNR aka specificity) and Positive Predictive Value (PPV aka precision) were recorded at the end of each round. This procedure was repeated 10 times with different random partitioning of the dataset resulting in 100 rounds of train-test process. Finally, the averages of the recorded classification scores were considered as the final cross-validated scores of the classifier.

The 10-times repetition of the CV procedure reduces the effect of bias on classification scores due to a relatively small number of samples. It ensures that the classification scores and hyperparameter tuning are not affected by any specific train-test partitioning of the dataset and the classifier is generalizable to future unseen samples.

### *F*_*0*.*5*_ score

Regarding ethical aspects of this study, we decided to minimize false positives, in the first place, while detecting ASD samples as much as possible. This goal could be achieved by maximization of PPV but it would be at a cost of decreasing TPR. To treat this issue, we chose to maximize *F*_*0*.*5*_ score which balances the PPV and the TPR while puts higher weight on PPV i.e. it pays more attention on minimization of false positives:

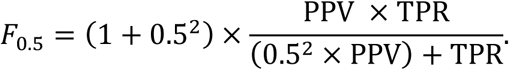

### Important feature extraction

An important goal of ML analysis in this study was to extract features that separate effectively NT babies from ASD ones. Our approach to this goal was 2-folded: 1) finding features that appears more frequently in the CV process; 2) finding features with the highest impact on the classifier’s decisions. In the first approach, selected features by the classifier in each CV fold were recorded. At the end of the CV process, the frequency of each feature was computed. Those features that appeared more frequently in the CV process were more important for classification.

In the second approach, the classifier was trained by all 252 samples and the classifier’s output was explained by the novel SHapley Additive exPlanations (SHAP) framework [75]. This method works in the level of each sample and feature and provides more details than the first approach. It provides SHAP values *s*_*ij*_ that indicates the impact of feature *j* on the classifier’s decision for child *i*. A positive or negative *s*_*ij*_ means that feature *j* pushes the classifier to classify the child *i* in the ASD or NT group, respectively. The higher the absolute value |*s*_*ij*_|, the bigger impact of feature *j* on the classifier’s decision for child *i*. On the other hand, *s*_*ij*_ ≈ 0 implies a very low impact.

The total impact of all features for all 252 children is computed as:

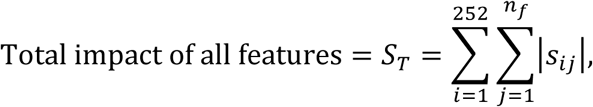

where *n*_*f*_ is the number of selected features by the FSS. The absolute impact of each feature is calculated as:

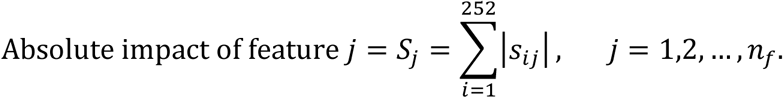

The relative impact of each feature in percent is given by:

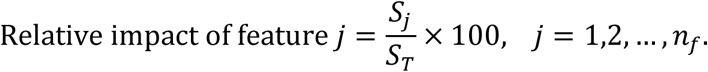

The relative impact is used to rank features and extract the most impactful ones.

### Statistical analysis

The difference in distribution of 116 collected features between NT and ASD groups were investigated by using conventional statistical hypothesis tests. For categorical features, Chi-squared test (*χ* ^2^) was used. The two-sided Welch’s t-test was applied on numerical parameters when the normality assumption was plausible according to the Shapiro-Wilk normality test. Otherwise, the nonparametric two-sided Mann-Whitney U test (MWU) was used. Moreover, the Benjamini–Hochberg procedure was employed to decrease the false discovery rate which adjusted the significance level to α = 0.001.

We used Analysis of covariance (ANCOVA) to model the fetal brain developmental trajectories, measured as cephalic perimeter (CP) from ultrasound acquisition during the 2^nd^ and 3^rd^ trimesters and before birth. It also let us to test if the growth rate (slope of regression line) changes between NT and ASD at each age (level of significance α = 0.05). Inspired by the collective CP distribution from the 2^nd^ trimester to before birth, a quadratic mixed effect model was fitted to determine brain growth rate in this period. Interaction term between ultrasound acquisition day and ASD or NT condition was considered as fixed effects. Random intercepts and slopes were included to take inter-individual baseline and growth rate variabilities into account.

Distributions of ASD CP percentile before birth revealed a negative skewness where about 38% of ASD children had large CP percentile (>90%). We conjectured that those fetuses have already had large CP in the 2^nd^ or 3^rd^ trimester. To examine that, those fetuses were separated from other ASD samples and they formed the “Large CPs ASD” group. The difference in CP percentile distributions of the 3 groups i.e. NT, ASD and Large CPs ASD, was checked by ANOVA with Tukey’s post-hoc test (when the normality assumption was plausible) or Kruskal-Wallis test with Dunn’s post-hoc test in the 2^nd^ and 3^rd^ trimesters and before birth. Moreover, the brain growth of this group was compared to that of other ASD and NT babies by using ANCOVA and a quadratic mixed effect model as described before.

### Implementation

All the programming and implementation of XGBoost was done on Python v.3.6 using NumPy v.1.18.1, Pandas v.0.24.2, scikit-learn v.0.22.1, Matplotlib v.3.0.2 and XGBoost v.0.80 libraries. Impact of parameters was calculated thanks to the SHAP library v.0.28.5. Moreover, we used the SciPy v.1.1.0 and StatsModels v.0.10.0 libraries for statistical tests, linear regression analysis and mixed effect analysis.

