## Supplementary figures for "Pregnancy data enable identification of relevant biomarkers and a partial prognosis of autism at birth"

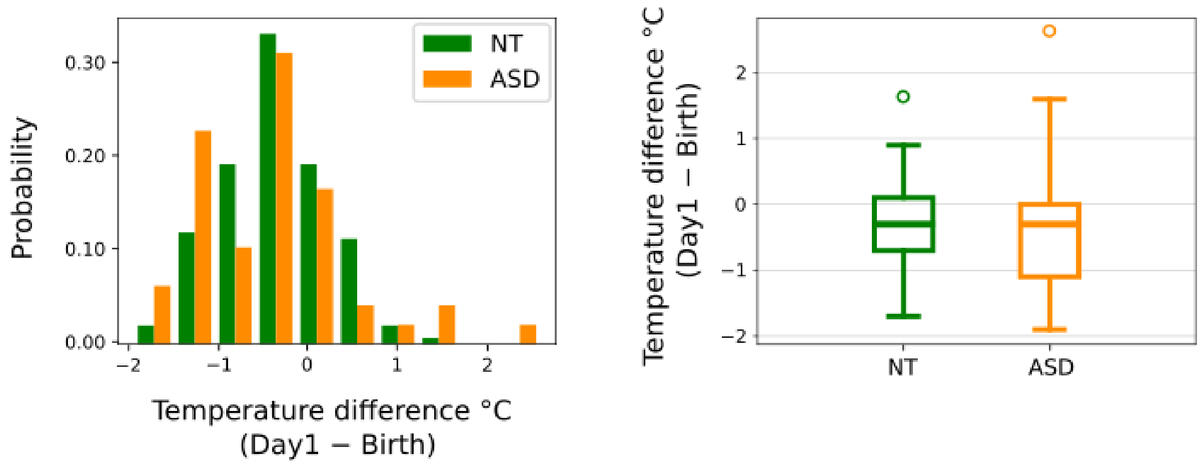


**Figure S1.** Left: Distribution of child's temperature difference between day 1 and birth in the Neurotypical (NT; green) and ASD (orange) groups. Right**:** Heterogeneity of temperature difference in the ASD group leads to a significant difference between the NT and ASD groups in two tails of the distributions such that the number of children with temperature difference more than 1°C is significantly larger in the ASD than in the NT group. For quantitative information, see Table 3.

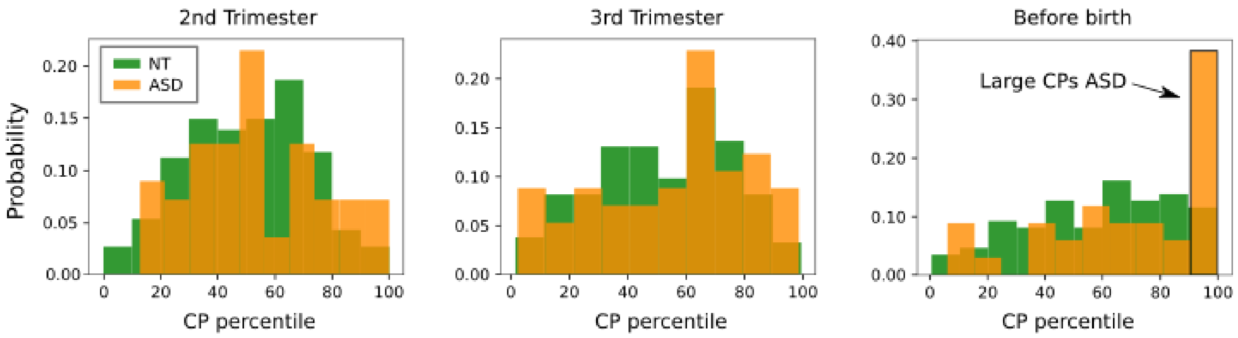


**Figure S2.** Distribution of cephalic perimeter (CP) percentiles in the NT (green) and ASD (orange) groups. During each 2nd and 3rd trimesters, the distribution of CP percentiles is quite similar in both groups. In contrast, there is a subpopulation of the ASD group with obviously large CP shortly before birth. We call it "Large CPs ASD" and analyzed them separately (see Fig. 4c and 4d).
