## Supplementary Table 2 for "Pregnancy data enable identification of relevant biomarkers and a partial prognosis of autism at birth"

**Supplementary Table S2**

For each feature, statistical information of sample distributions in ASD and Neurotypic (NT) groups are given. In case of categorical features, frequency (%) and number (n) of samples in each category are shown. In the case of numerical features, median, mean, standard error of mean, 95% confidence interval of mean are given respectively. Results of the statistical tests between ASD and NT groups are presented. Chi-2, Chi-squared test. T-test, Welch's t-test. MWU, Mann-Whitney U test.

| Feature | | Statistics | | | |
|---|---|---|---|---|---|
| | | ASD group | NT group | Stat. test | p-value |
| Sex | Male % (n) | 80.95 (51) | 48.70 (92) | Chi-2 | <0.001 |
| | Female % (n) | 19.05 (12) | 51.30 (97) | | |
| | Ratio M/F | 4.25 | 0.95 | | |
| Fetal rotation on head (days) | | 166 (178.35, 4.81, 168.72-187.98) | 180 (186.55, 2.30, 182.01-191.09) | MWU | 0.03 |
| Child's temperature difference day 1 − birth | > 1° % (n) | 41.66 (20) | 14.00 (21) | Chi-2 | <0.001 |
| | < 1° % (n) | 58.33 (28) | 86.00 (129) | | |
| Nasal bone length in T2 | | 7.50 (7.16, 0.20, 6.76-7.56) | 7.75 (7.68, 0.07, 7.55-7.81) | T-test | 0.017 |
| Foot length in T2 | | 41.00 (41.28, 0.55, 40.14-42.42) | 40.25 (40.26, 0.21, 39.84-40.69) | MWU | 0.10 |
| Femur length percentile in T3 | | 45.62 (45.60, 3.61, 38.36-52.84) | 38.20 (39.19, 1.63, 35.97-42.42) | MWU | 0.13 |

| Feature | | Statistics | | | |
|---|---|---|---|---|---|
| | | ASD group | NT group | Stat. test | p-value |
| Nasal bone length in T3 | | 11.00 (10.89, 0.20, 10.47-11.31) | 11.50(11.13, 0.11, 10.92-11.35) | MWU | 0.13 |
| Abdominal perimeter percentile in T2 | | 58.91 (59.02, 2.82, 53.36-64.68) | 55.38 (56.18, 1.67, 52.89-59.48) | MWU | 0.40 |
| Cephalic perimeter percentile in T2 | | 51.60 (54.61, 3.09, 48.42-60.79) | 51.60 (50.15, 1.54, 47.12-53.18) | T-test | 0.20 |
| Familial maternal history of auto-immune diseases | Yes % (n) | 19.05 (12) | 6.35 (12) | Chi-2 | 0.0064 |
| | No % (n) | 80.95 (51) | 93.65 (177) | | |
| White cells in T3 | | 9800.00 (10039.84, 403.10, 9233.52-10846.15) | 10230.00 (10494.00, 204.56, 10090.17-10897.83) | MWU | 0.12 |
| Real first trimester markers | | 4500.00 (10320.49, 2425.53, 5438.15-15202.84) | 7901.00 (15034.73, 2456.60, 10183.42-19886.05) | MWU | 0.11 |
| Cephalic perimeter percentile before birth | | 73.58 (69.03, 5.04, 58.78-79.28) | 62.56 (58.53, 2.74, 53.07-63.98) | MWU | 0.021 |
| pCO2 on arterial blood at the umbilical cord | | 56.05 (54.54, 1.60, 51.28-57.79) | 50.85 (50.43, 0.70, 49.05-51.81) | MWU | 0.028 |
| CMV | Immunized % (n) | 76.92 (20) | 36.62 (26) | Chi-2 | <0.001 |
| | Negative % (n) | 23.08 (6) | 63.38 (45) | | |
| Delta weight percentile between birth and day 1 | | 7.46 (10.51, 1.35, 7.80-13.23) | 6.79 (9.51, 0.68, 8.16-10.86) | MWU | 0.71 |
| Nucal transluciency percentile in T1 | | 25.00 (31.80, 3.93, 23.88-39.72) | 30.00 (36.63, 2.04, 32.62-40.65) | MWU | 0.34 |
| Fetal weight estimation percentile in T2 | | 55.75 (56.87, 2.87, 51.12-62.62) | 51.60 (51.36, 1.86, 47.69-55.03) | MWU | 0.18 |

| Feature | | Statistics | | | |
|---|---|---|---|---|---|
| | | ASD group | NT group | Stat. test | p-value |
| Lateral ventricle size percentile in T2 | | 5.00 (22.36, 5.58, 10.84-33.88) | 5.00 (16.53, 1.69, 13.19-19.87) | MWU | 0.78 |
| Duration of the first part of the labour (minute) | | 28.00 (249.74, 27.05, 195.65-303.84) | 267.00 (262.53, 15.00, 232.95-292.12) | MWU | 0.52 |
| Child's temperature at day 1 | | 36.70 (36.74, 0.05, 36.64-36.84) | 36.90 (36.91, 0.03, 36.85-36.96) | MWU | 0.0030 |
| Transverse diameter of the cerebellum percentile in T3 | | 57.50 (60.97, 4.48, 51.80-70.14) | 75.00 (70.82, 1.47, 67.92-73.72) | MWU | 0.029 |
| Child's temperature at birth | | 37.00 (37.09, 0.10, 36.88-37.30) | 37.20 (37.22, 0.04, 37.14-37.29) | T-test | 0.26 |
| Ratio IgM CMV | | 0.31 (0.30, 0.02, 0.26-0.34) | 0.26 (0.28, 0.02, 0.24-0.32) | MWU | 0.25 |
| IgG CMV | | 9950.00 (10913.64, 2034.00, 6683.70-15143.57) | 0.00 (4675.00, 963.19, 2750.23-6599.77) | MWU | < 0.001 |
| Fetal heart rate during labour (FIGO classification) | Pathological % (n) | 28.89 (13) | 10.20 (15) | Chi-2 | < 0.001 |
| | Suspect % (n) | 8.89 (4) | 28.57 (42) | | |
| | Normal % (n) | 62.22 (28) | 61.22 (90) | | |
| Apgar score at1minute | | 10.00 (8.73, 0.29, 8.15-9.31) | 10.00 (9.32, 0.11, 9.09-9.54) | MWU | 0.14 |
| Antibiotics | Yes % (n) | 19.35 (12) | 5.62 (10) | Chi-2 | 0.003 |
| | No % (n) | 80.65 (50) | 94.38 (168) | | |
| Real PAPP_A | | 10.04 (19.35, 4.08, 10.90-27.79) | 4.62 (8.76, 1.13, 6.52-11.01) | MWU | 0.004 |
| Ratio of cephalic perimeter to femoral length in T2 | | 5.04 (5.07, 0.04, 4.99-5.16) | 5.02 (5.06, 0.03, 5.01-5.11) | MWU | 0.99 |

| Feature | | Statistics | | | |
|---|---|---|---|---|---|
| | | ASD group | NT group | Stat. test | p-value |
| Duration of the rupture of the membranes (minutes) | | 147.00 (254.22, 41.73, 170.80-337.64) | 222.00 (401.04, 61.59, 279.53-522.54) | MWU | 0.20 |
| Biparietal diameter percentile in T2 | | 49.80 (47.58, 3.08, 41.41-53.75) | 42.85 (43.42, 1.75, 39.97-46.88) | MWU | 0.28 |
| Craniocaudal length percentile in T1 | | 80.00 (82.72, 1.38, 79.91-85.54) | 80.00 (78.92, 1.25, 76.45-81.39) | MWU | 0.27 |
| Origin | African % (n) | 22.22 (14) | 10.05 (19) | Chi-2 | 0.010 |
| | Caucasians % (n) | 61.90 (39) | 80.42 (152) | | |
| | Other % (n) | 15.87 (10) | 9.52 (18) | | |
| Real FBhCG | | 30.55 (45.71, 7.63, 29.93-61.49) | 29.62 (36.76, 2.82, 31.17-42.36) | MWU | 0.46 |
| Ombilical doppler in T3 | | 0.64 (0.63, 0.01, 0.61-0.66) | 0.63 (0.63, 0.01, 0.62-0.64) | T-test | 0.87 |
| PAPP_A (DoE) | | 0.85 (0.83, 0.22, 0.38-1.27) | 0.13 (0.18, 0.08, 0.02-0.33) | MWU | 0.005 |
| Ratio of biparietal diameter to cranio-caudal length in T1 | | 0.34 (0.34, 0.01, 0.33-0.36) | 0.34 (0.35, 0.003, 0.34-0.35) | MWU | 0.54 |
| Biparietal diameter percentile in T3 | | 33.71 (38.47, 4.12, 30.21-46.73) | 35.55 (37.41, 1.90, 33.67-41.15) | MWU | 0.86 |
| FBhCG (DoE) | | 0.47 (0.46, 0.18, 0.10-0.83) | -0.16 (0.06, 0.10, -0.14-0.25) | MWU | 0.011 |
| Fibrinogen in T3 | | 4.85 (4.88, 0.11, 4.66-5.11) | 4.81 (4.84, 0.07, 4.71-4.98) | MWU | 0.93 |
| Platelets T3+DS1 | | 241.00 (251.03, 8.91, 233.20-268.86) × $10^3$ | 245.00 (247.87, 4.74, 238.52-257.23) × $10^3$ | MWU | 0.78 |

| Feature | Statistics | | | |
|---|---|---|---|---|
| | ASD group | NT group | Stat. test | p-value |
| Transverse diameter of the cerebellum percentile in T2 | 55.00 (50.81, 4.11, 42.41-59.20) | 55.00 (56.59, 1.50, 53.64-59.55) | MWU | 0.32 |
| Number of miscarriages | 0.00 (0.24, 0.08, 0.08-0.39) | 0.00 (0.46, 0.07, 0.33-0.58) | MWU | 0.044 |
| Ratio of cephalic perimeter to femoral length in T3 | 4.74 (4.73, 0.04, 4.65-4.80) | 4.76 (4.77, 0.02, 4.74-4.81) | MWU | 0.26 |
| First mechonium emission (minute) | 395.00 (439.22, 54.70, 329.35-549.08) | 377.50 (431.80, 34.08, 364.48-499.12) | MWU | 0.74 |
| Abdominal perimeter percentile in T3 | 50.80 (53.65, 3.53, 46.58-60.72) | 53.19 (52.16, 1.77, 48.66-55.65) | MWU | 0.71 |
| Mother's body mass index before pregnancy | 23.40 (25.19, 0.84, 23.51-26.87) | 22.90 (24.41, 0.44, 23.55-25.27) | MWU | 0.44 |
| Weight percentile at birth | 38.76 (42.39, 3.77, 34.86-49.92) | 39.66 (42.25, 1.99, 38.33-46.17) | MWU | 0.97 |
| Folic Acid | 0.40 (0.58, 0.16, 0.27-0.90) | 0.40 (0.47, 0.05, 0.36-0.57) | MWU | 0.048 |
| Weight percentile at Day 1 | 22.86 (33.85, 3.98, 25.86-41.84) | 29.66 (34.81, 1.99, 30.88-38.74) | MWU | 0.57 |
| Ratio of femoral length to cranio-caudal length in T1 | 0.12 (0.13, 0.01, 0.12-0.14) | 0.13 (0.13, 0.002, 0.12-0.13) | MWU | 0.50 |
| Hemoglobin in T3 | 11.60 (11.75, 0.14, 11.47-12.03) | 11.80 (11.87, 0.09, 11.69-12.05) | T-test | 0.48 |
| Ratio cephalic perimeter to femoral length before birth | 4.64 (4.63, 0.05, 4.54-4.72) | 4.57 (4.61, 0.03, 4.54-4.67) | MWU | 0.56 |
| Ratio of cephalic perimeter to height | 0.70 (0.70, 0.00, 0.69-0.71) | 0.70 (0.70, 0.00, 0.69-0.70) | MWU | 0.35 |
| Abdominal perimeter percentile before birth | 67.86 (59.19, 4.82, 49.38-69.00) | 53.59 (52.93, 3.08, 46.80-59.05) | MWU | 0.28 |

| Feature | | Statistics | | | |
|---|---|---|---|---|---|
| | | ASD group | NT group | Stat. test | p-value |
| Term of birth (days) | | 274.00 (271.51, 2.28, 266.94-276.07) | 274.00 (271.06, 1.07, 268.94-273.18) | MWU | 0.46 |
| Study level | Primary % (n) | 20.63 (13) | 6.35 (12) | Chi-2 | 0.004 |
| | Secondary % (n) | 53.97 (34) | 60.85 (115) | | |
| | Superior % (n) | 25.40 (16) | 32.80 (62) | | |
| Umbilical doppler before birth | | 0.59 (0.58, 0.01, 0.55-0.60) | 0.59 (0.60, 0.01, 0.59-0.62) | T-test | 0.098 |
| Biparietal diameter percentile in T1 | | 50.00 (54.18, 4.44, 45.14-63.22) | 50.00 (54.55, 1.62, 51.36-57.75) | MWU | 0.94 |
| Maternal tobacco use | | 0.00 (1.37, 0.58, 0.21-2.53) | 0.00 (3.06, 0.53, 2.01-4.10) | MWU | 0.019 |
| Cephalic perimeter percentile in T3 | | 61.04 (55.36, 3.47, 48.41-62.32) | 55.38 (52.23, 1.70, 48.87-55.59) | MWU | 0.31 |
| Fetal weight estimation percentile in T3 | | 36.68 (43.42, 4.14, 35.13-51.71) | 32.98 (37.11, 1.80, 33.55-40.67) | MWU | 0.26 |
| Breastfeeding | Artificial % (n) | 17.86 (10) | 30.10 (56) | Chi-2 | 0.009 |
| | Maternal % (n) | 64.29 (36) | 63.98 (119) | | |
| | Mixed % (n) | 17.86 (10) | 5.91 (11) | | |
| Cephalic perimeter percentile in day 1 | | 47.13 (49.07, 3.58, 41.89-56.24) | 45.62 (47.73, 1.79, 44.20-51.26) | MWU | 0.83 |
| Femur length percentile in T2 | | 53.30 (54.85, 2.97, 48.89-60.81) | 51.60 (51.52, 1.62, 48.32-54.73) | MWU | 0.40 |
| Number of Caesarean sections | | 0.00 (0.19, 0.06, 0.06-0.32) | 0.00 (0.14, 0.03, 0.07-0.20) | MWU | 0.41 |

| Feature | | Statistics | | | |
|---|---|---|---|---|---|
| | | ASD group | NT group | Stat. test | p-value |
| Duration of the second part of the labour (minute) | | 14.00 (39.54, 6.88, 25.80-53.28) | 20.00 (55.68, 5.63, 44.58-66.79) | MWU | 0.31 |
| Vitamin D | | 0.00 (29.03, 5.81, 17.41-40.65) × $10^3$ | 0.00 (25.28, 3.27, 18.83-31.73) × $10^3$ | MWU | 0.57 |
| Duration of epidural analgesia (minute) | | 207.00 (205.02, 21.55, 161.93-248.10) | 210.00 (217.29, 14.28, 189.13-245.46) | MWU | 0.85 |
| Apgar score at 3 minutes | | 10.00 (9.44, 0.20, 9.05-9.84) | 10.00 (9.69, 0.07, 9.54-9.83) | MWU | 0.17 |
| First urine (minute) | | 537.50 (555.35, 75.97, 402.82-707.87) | 517.50 (610.65, 43.90, 523.94-697.37) | MWU | 0.45 |
| Pregnancy weight gain | | 13.00 (12.75, 0.74, 11.26-14.25) | 13.00 (12.91, 0.42, 12.08-13.74) | T-test | 0.86 |
| Maternal history of infectious diseases | Yes % (n) | 20.63 (13) | 10.58 (20) | Chi-2 | 0.067 |
| | No % (n) | 79.37 (50) | 89.41 (169) | | |
| Number of urgent Caesarean sections | | 0.00 (0.16, 0.06, 0.05-0.27) | 0.00 (0.10, 0.02, 0.05-0.14) | MWU | 0.43 |
| Skin to skin protocol | Immediate % (n) | 20.63 (13) | 32.45 (61) | Chi-2 | 0.20 |
| | Precoce % (n) | 19.05 (12) | 15.43 (29) | | |
| | No % (n) | 60.32 (38) | 52.13 (98) | | |
| Oxytocin during labor | | 0.00 (7.73, 1.62, 4.49-10.97) | 0.00 (8.22, 0.94, 6.37-10.07) | MWU | 0.59 |
| Fetal cardiac short term variability before birth | | 10.30 (11.84, 0.95, 9.89-13.80) | 10.90 (11.35, 0.35, 10.66-12.03) | MWU | 0.99 |
| Femur length percentile before birth | | 32.81 (38.24, 4.59, 28.90-47.58) | 28.42 (33.57, 2.97, 27.67-39.47) | MWU | 0.35 |

| Feature | | Statistics | | | |
|---|---|---|---|---|---|
| | | ASD group | NT group | Stat. test | p-value |
| Familal maternal history of endocrine diseases | Yes % (n) | 41.27 (26) | 38.62 (73) | Chi-2 | 0.82 |
| | No % (n) | 58.73 (37) | 61.38 (116) | | |
| Fetal weight estimation percentile before birth | | 89.79 (71.35, 5.80, 59.54-83.15) | 77.34 (62.11, 4.05, 54.07-70.16) | MWU | 0.17 |
| Instrumental delivery | VB % (n) | 66.67 (42) | 63.49 (120) | Chi-2 | 0.89 |
| | Cesar % (n) | 23.81 (15) | 24.87 (47) | | |
| | Forceps % (n) | 4.76 (3) | 7.41 (14) | | |
| | Ventouse % (n) | 4.76 (3) | 4.23 (8) | | |
| Serology of toxoplasmaosisa | Immunized % (n) | 31.75 (20) | 34.92 (66) | Chi-2 | 0.76 |
| | Negative % (n) | 68.25 (43) | 65.08 (123) | | |
| Placenta Grannum classification in T3 | | 1.00 (0.97, 0.13, 0.70-1.24) | 1.00 (0.94, 0.05, 0.84-1.04) | MWU | 0.79 |
| Biparietal diameter percentile before birth | | 38.20 (48.03, 5.28, 37.29-58.77) | 33.71 (38.77, 3.11, 32.59-44.95) | MWU | 0.13 |
| Femur length percentile in T1 | | 50.00 (49.74, 4.08, 41.41-58.07) | 50.00 (51.52, 1.44, 48.68-54.36) | MWU | 0.61 |
| Glycemia at birth | | 0.65 (0.66, 0.03, 0.59-0.72) | 0.63 (0.66, 0.02, 0.62-0.69) | T-test | 0.99 |
| Number of scheduled Caesarean sections | | 0.00 (0.03, 0.02, -0.01-0.08) | 0.00 (0.04, 0.02, 0.01-0.08) | MWU | 0.99 |
| Fetal cardiac rhythm | Normal % (n) | 62.22 (28) | 61.22 (90) | Chi-2 | 0.96 |

| Feature | | Statistics | | | |
|---|---|---|---|---|---|
| | | ASD group | NT group | Stat. test | p-value |
| FIGO 2 groups | Pathological % (n) | 37.78 (17) | 38.78 (57) | | |
| Foot length in T3 | | 67.00 (69.40, 1.51, 65.97-72.83) | 65.50 (65.39, 1.11, 63.11-67.68) | T-test | 0.046 |
| Pathological pH | Normal % (n) | 82.46 (47) | 90.76 (167) | Chi-2 | 0.13 |
| | Pathological % (n) | 17.54 (10) | 9.24 (17) | | |
| Maternal history of endocrine diseases | Yes % (n) | 14.29 (9) | 12.70 (24) | Chi-2 | 0.91 |
| | No % (n) | 85.71 (54) | 87.30 (165) | | |
| Familial paternal history of endocrine diseases | Yes % (n) | 10.34 (6) | 15.30 (28) | Chi-2 | 0.47 |
| | No % (n) | 89.66 (52) | 84.70 (155) | | |
| Controlled diabetes | Yes % (n) | 10.71 (6) | 14.86 (26) | Chi-2 | 0.73 |
| | No % (n) | 8.93 (5) | 8.00 (14) | | |
| | NS % (n) | 80.36 (45) | 77.14 (135) | | |
| Size of lateral ventricle percentile in T3 | | 3.00 (12.10, 4.54, 2.71-21.50) | 3.00 (6.81, 1.19, 4.45-9.18) | MWU | 0.33 |
| Antibiotics during labour | Yes % (n) | 15.87 (10) | 22.75 (146) | Chi-2 | 0.33 |
| | No % (n) | 84.13 (53) | 77.25 (43) | | |
| Apgar score at 5 minutes | | 10.00 (9.67, 0.16, 9.35-9.98) | 10.00 (9.86, 0.06, 9.74-9.97) | MWU | 0.37 |

| Feature | | Statistics | | | |
|---|---|---|---|---|---|
| | | ASD group | NT group | Stat. test | p-value |
| Hearing test symmetry | Symmetric % (n) | 91.49 (43) | 93.39 (113) | Chi-2 | 0.92 |
| | Asymmetric % (n) | 8.51 (4) | 6.61 (8) | | |
| Hearing test | Normal % (n) | 78.26 (36) | 80.17 (97) | Chi-2 | 0.95 |
| | Abnormal % (n) | 21.74 (10) | 19.83 (24) | | |
| Treatment for labor induction | Yes % (n) | 20.63 (13) | 22.75 (43) | Chi-2 | 0.86 |
| | No % (n) | 70.37 (50) | 77.25 (146) | | |
| Type of delivery | Vaginal % (n) | 69.84 (44) | 73.02 (138) | Chi-2 | 0.75 |
| | Caesarean % (n) | 30.16 (19) | 26.98 (51) | | |
| Coagulation in T3 | Normal % (n) | 89.83 (53) | 88.34 (144) | Chi-2 | 0.95 |
| | Abnormal % (n) | 10.17 (6) | 11.66 (19) | | |
| Streptococcus B vaginal swab | Positive % (n) | 10.53 (6) | 7.22 (13) | Chi-2 | 0.60 |
| | Negative %(n) | 89.47 (51) | 92.78 (167) | | |
| Apgar score at10 minutes | | 10.00 (9.78, 0.11, 9.55-10.00) | 10.00 (9.91, 0.04, 9.83-9.99) | MWU | 0.074 |
| Maternal history of auto-immune diseases | Yes % (n) | 23.81 (15) | 19.05 (36) | Chi-2 | 0.53 |
| | No % (n) | 76.19 (48) | 80.95 (153) | | |
| Gestational diabetus | Yes % (n) | 16.07 (9) | 19.43 (34) | Chi-2 | 0.72 |

| Feature | | Statistics | | | |
|---|---|---|---|---|---|
| | | ASD group | NT group | Stat. test | p-value |
| | No % (n) | 83.93 (47) | 80.57 (141) | | |
| Aspegic | Yes % (n) | 1.61 (1) | 8.99 (16) | Chi-2 | 0.096 |
| | No % (n) | 98.39 (61) | 91.01 (162) | | |
| Rh blood group system | Positive % (n) | 84.13 (53) | 85.19 (161) | Chi-2 | 1.00 |
| | Negative % (n) | 15.87 (10) | 14.81 (28) | | |
| IgM | Positive % (n) | 4.55 (1) | 0 (0) | Chi-2 | 0.59 |
| | Negative % (n) | 95.45 (21) | 100 (61) | | |
| Corticosteroids | Yes % (n) | 6.45 (4) | 7.30 (13) | Chi-2 | 0.95 |
| | No % (n) | 93.55 (58) | 92.70 (165) | | |
| Familial paternal history of auto-immune diseases | Yes % (n) | 6.90 (4) | 4.37 (8) | Chi-2 | 0.67 |
| | No % (n) | 93.10 (54) | 95.63 (175) | | |
| Paternal history of endocrine diseases | Yes % (n) | 0 (0) | 0.55 (1) | Chi-2 | 0.54 |
| | No % (n) | 100 (58) | 99.45 (182) | | |
| Paternal history of infectious diseases | Yes % (n) | 1.72 (1) | 0 (0) | Chi-2 | 0.54 |
| | No % (n) | 98.28 (57) | 100 (183) | | |
| Paternal history of | Yes % (n) | 3.45 (2) | 2.19 (4) | Chi-2 | 0.96 |

| Feature | | Statistics | | | |
|---|---|---|---|---|---|
| | | ASD group | NT group | Stat. test | p-value |
| auto-immune diseases | No % (n) | 96.55 (56) | 97.81 (179) | | |
| Medically assisted procreation | Yes % (n) | 6.35 (4) | 12.70 (24) | Chi-2 | 0.25 |
| | No % (n) | 93.65 (59) | 87.30 (165) | | |
| Serology of rubella | Immunized % (n) | 92.06 (58) | 94.18 (178) | Chi-2 | 0.76 |
| | Negative % (n) | 7.94 (5) | 5.82 (11) | | |
