## Supplementary Table 3 for "Pregnancy data enable identification of relevant biomarkers and a partial prognosis of autism at birth"

**Supplementary Table S3**

Complete list of all features collected from the 1st, 2nd and 3rd trimester, before birth, and during the post-natal period.

| List of collected features | | |
|---|---|---|
| **General information about parents** | | |
| • Mother's age<br>• Father's age<br>• Family history of mother and father (endocrine and autoimmune)<br>• History of mother and father (endocrine, infectious, and autoimmune) | • Maternal body mass index before pregnancy<br>• Maternal serologies for rubella, toxoplasmosis and CMV,<br>• Maternal blood RH type | • Maternal tobacco use<br>• Ethnic origin<br>• Level of study<br>• Number of pregnancies<br>• Number of miscarriages<br>• Number of emergency or scheduled Caesarean sections |
| **Treatments during pregnancy** | | |
| • Vitamin D<br>• Folic acid<br>• Antibiotics | • Aspirin<br>• Corticosteroids | |
| **General pregnancy data** | | |
| • Real first trimester marker (Actual HT21 rate)<br>• Gestational diabetes<br>• Controlled gestational diabetes<br>• Weight gain during pregnancy | • PAPP-A real dosage and DoE<br>• FBhCG real dosage and DoE<br>• Term of fetal rotation on head | |
| **First trimester of pregnancy** | | |
| • Nuchal translucency measurement and percentile<br>• Biparietal diameter measurement and percentile<br>• Femur length measures and percentile | • Cranio-caudal length measurement and percentile<br>• Biparietal diameter to cranio-caudal length ratio<br>• Femur length measurement and percentile | • Femoral length to cranio-caudal ratio |
| **Second trimester of pregnancy** | | |
| • Biparietal diameter measurement and percentile<br>• Femur length measurement and percentile | • Cephalic perimeter measurement and percentile<br>• Ratio of cephalic perimeter to femoral length | • Transverse diameter of the cerebellum measurement and percentile<br>• Lateral ventricle measurement and percentile<br>• Nasal bone length<br>• Foot length |

| | | |
|---|---|---|
| • Abdominal perimeter measurement and percentile | • Estimation of fetal weight measurement and percentile | |
| **Third trimester of pregnancy** | | |
| • Biparietal diameter measurement and percentile<br>• Femur length measurement and percentile<br>• Abdominal perimeter measurement and percentile<br>• Umbilical Doppler | • Cephalic perimeter measurement and percentile<br>• Ratio of cephalic perimeter to femoral length<br>• Estimation of fetal weight measurement and percentile | • Transverse diameter of the cerebellum measurement and percentile<br>• Lateral ventricle measurement and percentile<br>• Nasal bone length<br>• Foot length<br>• Placenta Grannum classification |
| **Prepartum phase** | | |
| • Biparietal diameter measurement and percentile<br>• Femur length measurement and percentile<br>• Abdominal perimeter measurement and percentile<br>• Cephalic perimeter measurement and percentile | • Ratio of cephalic perimeter to femoral length<br>• Umbilical Doppler<br>• Short-term variability in fetal heart rate<br>• Estimation of fetal weight measurement and percentile | • Streptococcus B vaginal swab<br>• White cells<br>• Hemoglobin<br>• Platelets<br>• Coagulation with fibrinogenemia |
| **Birth** | | |
| • Term of birth<br>• Type of delivery<br>• Treatment for labor induction<br>• Antibiotic during labor<br>• FIGO stage of fetal heart rate during labor | • Duration of the first and second part of the labor<br>• Duration of the rupture of the membranes | • Duration of epidural analgesia<br>• Use of oxytocin in labor maintenance |
| **Child** | | |
| • Sex<br>• Apgar score at 1, 3, 5 and 10 minutes<br>• Birth weight and percentile<br>• Temperature of the child at birth<br>• Skin-to-skin protocol<br>• Glycemia at birth | • pH and pCO2 on arterial blood at the umbilical cord<br>• Weight and percentile at D1<br>• Delta weight and percentile between birth and D1<br>• Cephalic perimeter measures and percentile at day 1 | • Cephalic perimeter to size ratio<br>• Hearing test<br>• Type of breastfeeding<br>• Mechonium emission<br>• Emission of first urine<br>• Temperature at D1<br>• Delta temperature between birth and D1 |
